# Association Studies of Phage Metagenomes from Aquatic Environment Identify Known and Potentially New Key Players in the 2,6-Diaminopurine Biosynthesis Pathway

**DOI:** 10.1101/2025.10.22.683976

**Authors:** Weiwei Yang, Shuang-yong Xu, Nan Dai, Ivan R. Corrêa, Laurence Ettwiller

## Abstract

Advances in next-generation sequencing (NGS) and bioinformatics have expanded the potential of metagenomic studies, particularly for unculturable organisms such as phages with heavily modified genomes. In the evolutionary arms race between phages and their hosts, some phages develop a strategy to fully modify their DNAs to evade host-encoded antiphage defenses. Here, we developed a restriction-enzyme based method to selectively enrich all phages containing 2-aminoadenine (diaminopurine or dZ) directly from purified metagenomic DNA from environmental samples. Applying our previously established metagenomic genome-phenome association (metaGPA) pipeline, we identified 116 protein domains significantly associated with dZ phages. The top candidate domains fully recapitulated the phage-encoded dZ biosynthetic pathway. Study of individual enzymes, including PurZ and YfbR-like dATPase at residue resolution revealed key amino acids for the distinction between dZ and dA substrate. Additionally, we identified and validated two novel DNA polymerases with enhanced dZ incorporation activity and uncovered nucleoside 2-deoxyribosyltransferase (NDT) as a potential new component of the dZ biosynthetic pathway. Collectively, these findings refine current models of the dZ biosynthesis pathway, potentially reveal previously unrecognized enzymatic components, and expand the understanding of phage biology with implications for synthetic biology and DNA engineering.

## Introduction

Bacteria and bacteriophages have co-evolved for billions of years, engaging in a continuous molecular arms race. This evolutionary pressure has driven the development of an extensive repertoire of antiphage defense systems in bacteria, including restriction-modification systems, CRISPR-Cas, and abortive infection mechanisms. In response, phages have evolved diverse countermeasures to circumvent these defenses. One such strategy involves the incorporation of chemically modified bases into their genomes (1), enabling them to evade host defense systems that rely on the recognition and cleavage of foreign DNA, such as restriction endonucleases (REases).

As these phages may encounter endonucleases with various sequence specificities, bases are partially or fully modified masking their cognate canonical counterparts. One such modification termed dZ also known as 2,6-diaminopurine has been found in various phages (2–5). The core biosynthetic pathway for dZ formation has recently been characterized. dZ genome phages encode a complex Z-cluster made of three major genes: a homologue of adenylosuccinate synthetase PurA (PurZ), a dATP triphosphohydrolase (DatZ) and a dGTP/dATP diphosphohydrolase. PurZ catalyzes the condensation from dGMP and aspartate to form aminodeoxyadenylosuccinate (ADAS) deoxynucleotide monophosphate, which is followed by stepwise reactions to generate dZTP catalyzed by host encoded PurB and GMP kinase (3, 5, 6). A recent study characterizes PurZ0, suggesting the progressive evolution from archaeal PurA to phage PurZ (7). The transition to use ATP and dATP as the phosphate donor of PurZ is a result of adaptation to ZTCG-genome by depleting dATP in the nucleotide pool. DatZ and dGTP/dATP diphosphohydrolase are also part of the machinery to eliminate dATP from the available nucleotides (2, 5). The substrate of PurZ, dGMP, is supplied from dGTP/dATP diphosphohydrolase with exception of cyanophage S-2L, that encodes a dGTP-specific diphosphohydrolase (MazZ) (2). In the Wayne-like clades of phages, a homologue of deoxyuridine phosphatase (dUTPase) may play the role of DatZ or MazZ (8). To incorporate dZTP into the genome, dZ phages encode a viral DNA polymerase DpoZ that demonstrates greater catalytic efficiency with dZTP than with dATP (4). Cyanophage S-2L instead encodes a PrimPol family enzyme which doesn’t seem to have selectivity of A or Z; however, the dATP specific phosphatase DatZ is responsible for the absence of adenine in its genome (2, 4).

In addition to revealing a mechanism for the biosynthesis of a unique DNA modification, the study of dZ phages and their maintenance of the unusual ZTCG genome opens new avenues for research into alternative synthetic organisms, phage anti-defense systems, and noncanonical DNA polymerization. Furthermore, a distinctive chemical property of base Z—its ability to form stronger hydrogen bonds with thymine—has attracted interest for its potential in anti-degradation therapeutics and high-capacity DNA-based data storage materials. To explore the diversity of dZ phages and their biosynthetic pathways in native microbiomes, we developed an enzymatic method to selectively enrich dZ phages and dZ-containing DNA sequences from environmental samples. We then applied a metagenomic genome-phenome association (metaGPA) approach to identify protein domains associated with dZ phages and comprehensively characterized the core dZ biosynthetic pathway. Our study uncovered several novel dZ biosynthesis associated protein domains and provided residue-level insights into the refined functions of PurZ and dATPase (DatZ). Notably, we validated two novel DpoZs which significantly enhance in vivo incorporation of base Z into E. coli. Interestingly, our findings also suggest a potential additional player in dZ biosynthesis: nucleoside 2-deoxyribosyltransferase, which appears to exhibit nucleoside substrate specificity.

## Results

### 1. Development of an enzymatic method to enrich dZ containing-DNA

Environmental microbiomes provide rich reservoirs for studying phages with DNA containing modified bases. To enrich genomic DNA predicted to contain dZ directly from the environment, we designed an enzyme cocktail strategy composed of seven distinct REases that restrict canonical dA DNA but do not have activity (or have low activity) on dZ containing DNA. The selection of REases was based on our previous work characterizing subsets of REases for presence or absence of activity on dZ containing amplicons (9) (**Materials and Methods**). The REase cocktail targets a range of frequent 4-6 bp recognition motifs, ensuring that most fragments containing dA are digested, while dZ-containing DNA remains intact. The average cutting frequency of the enzyme cocktail for restricting dA-containing phages is estimated to be approximately 1 cut per 82 bp, assuming an equimolar composition of dA, dT, dC, and dG. For positive control, dZ containing DNA was synthesized *in vitro*. Following digestion, the relative amounts of dZ or dA remained were measured by qPCR. This experiment confirmed a 99.3% recovery of dZ DNA and up to 96.5% depletion of canonical dA DNA (**Fig. 1A**).

**Figure 1.**
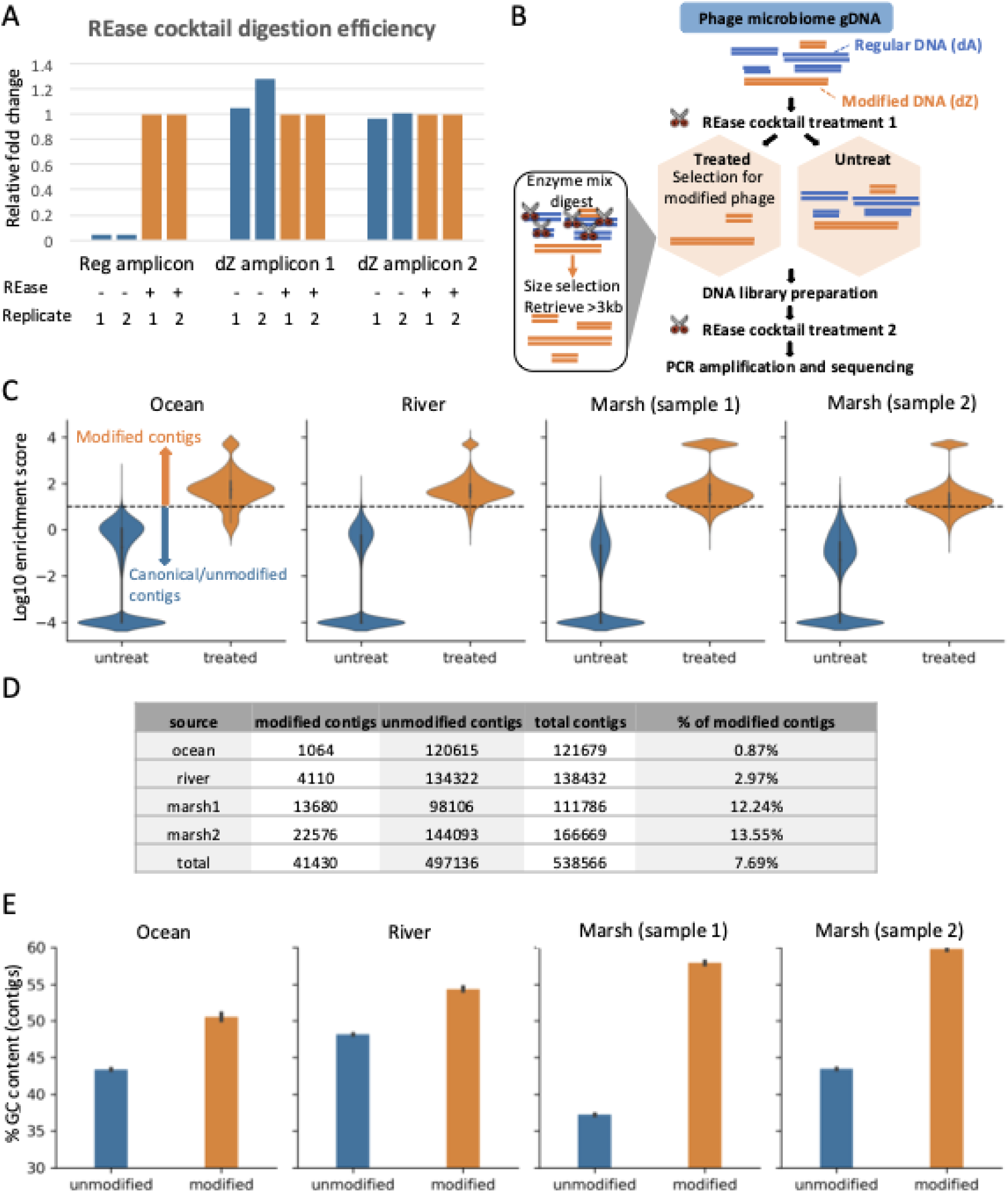
REase cocktail strategy efficiently enriches a subset of dZ-containing viral genomic DNA from aquatic samples. **A**) Digestion of canonical (left panel) and dZ containing amplicons (middle and right panels) using the REase cocktail.. dZ-DNA was synthesized by replacing dATP with dZTP in the PCR reaction. qPCR targeting canonical and dZ amplicons were conducted after REase cocktail treatment or using an untreated control to measure digestion efficiency. Cts were normalized to the corresponding untreated control for calculation of the relative fold change using delta delta Ct method. Digestion and qPCR were performed in replicates. **B**) Scheme of REase-based method to enrich and sequence predicted dZ-containing phage from viral fraction of four aquatic samples. **C**) Violin plot picturing the log10 enrichment scores of controls (blue) and treated (orange) contigs from four phage microbiome samples. An Enrichment Score (ES) cutoff of 10 was set to group contigs into dZ-modified contigs (ES > =10) and canonical dA contigs (ES <10). Contigs with infinite enrichment scores were capped and plotted at a value of 10000. **D**) Summary table of the number of assembled contigs (predicted dZ-modified or unmodified/canonical). **E**) GC content for predicted dZ-modified contigs (orange bars) and canonical/unmodified dA contigs (blue bars).

### 2. Application of the REase cocktail strategy to viral DNA from aquatic samples reveals dZ containing organisms

We applied the enzyme cocktail strategy to genomic DNA isolated from the viral fraction of the four aquatic samples comprising ocean, river, or salt marsh. To further reduce the background from undigested DNA, the enzyme treatment was performed twice, once before DNA fragmentation and once after adaptor ligation during library preparation. The remaining intact DNA fragments presumably containing the dZ modification were amplified and subjected to short-read sequencing using Illumina. An untreated control of the same viral fraction was performed in parallel for comparison (**Fig. 1B**).

High throughput sequencing resulted in between 10 - 50 million paired-end reads per sample. Reads were assembled into contigs using metaSPAdes (10). Assemblies from both control and treated sample pairs were combined, the redundancy was removed and the reads from the control/treated sample pair were mapped back to the combined assemblies using bowtie2. Using our previously established metagenomics genome-phenome association (metaGPA) pipeline (11), we assigned each *de novo* assembled contig with an enrichment score representing the fold difference in the normalized read counts between the treated samples and the untreated controls. Contigs that carry dZ modifications were expected to have high enrichment scores compared to the ones with canonical dA (**Fig. 1C**). Contigs with an enrichment score greater or equal to 10 were defined as exhibiting resistance to REase cocktail treatment and therefore predicted to contain the dZ modification (dZ modified contigs). We identified a total of 41,430 dZ-modified contigs out of a total of 497,136 contigs with size greater than 500 bp from the four environmental phage metagenome samples. In line with the fact that the majority of phage genomes are unmodified, this number corresponds to an average of ∼7.7% dZ-modified contigs (**Fig. 1D**). Contigs from salt marsh viriomes exhibited higher frequencies (12–13% of contigs) compared to that of ocean and river viriomes (0.8–3%), suggesting a greater prevalence of dZ genomic DNA in salt marsh environments. As reported in (12), we also observed higher GC-content in these modified contigs compared to that of canonical contigs (**Fig. 1E**) (12).

### 3. Association study using metaGPA framework identifies adenylosuccinate synthetase as the top domain associated with REase resistance

The metaGPA pipeline was applied to identify protein domains significantly associated with dZ modified contigs. Specifically, contigs were translated into all 6 frames and annotated using Pfam hidden markov models (HMMs). Fisher’s exact test was applied to each annotated protein domain to assess their likelihood of being enriched within the dZ modified contigs. A significant *p*-value indicates a non-random association between a given protein domain and the modified contigs, suggesting a potential role in dZ modification. metaGPA association analysis was performed independently on each of the four individual samples as well as on the combined dataset (see **Materials and Methods**).

In both the composite and three of the individual datasets, the adenylosuccinate synthetase domain (Pfam accession: PF00709) was found as the top associated protein domain (**Fig. 2A** and **Supplementary Fig. S1A-D**). This HMM representing the superfamily of adenylosuccinate synthetases includes profiles of the canonical adenylosuccinate synthetase (PurA), which is an enzyme that plays an important role in A base synthesis (13). It also includes the PurZ enzyme family (3, 5, 6) shown to catalyze the conversion of 2’-deoxyguanosine 5’-monophosphate (dGMP) into 2-amino-2’-deoxyadenosine 5’-monophosphate (dZMP), a key step of the dZ biosynthetic pathway (3, 5) in dZ phages (e.g. *cyanophage* S-2L) (3, 5).

**Figure 2.**
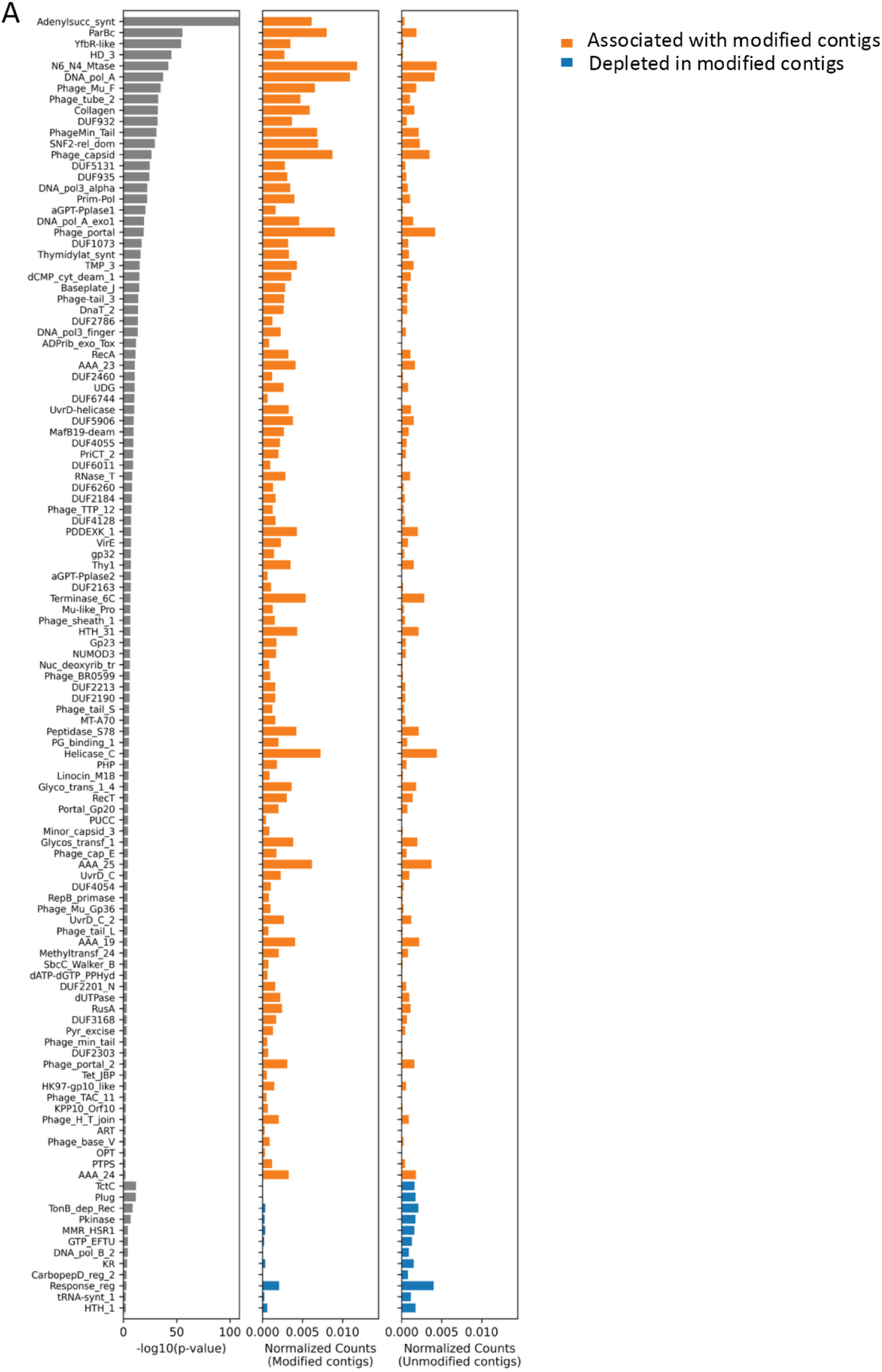
metaGPA identifies protein domains associated with dZ modification and recapitulates dZ biosynthetic pathways. **A**) Top 116 domains significantly (two-tailed p value < 0.01) associated with the dZ modification (orange bars) and 12 domains significantly (two-tailed p value < 0.01) depleted in the modified contigs. Left: p value of each Pfam association analysis; middle: normalized counts in metaGPA predicted dZ-modified contigs; right: normalized counts in unmodified contigs. Enriched Pfams in modified contigs are marked in orange and depleted Pfams in modified contigs are marked in blue.

To validate whether metaGPA specifically associates PurZ with dZ-modified contigs, we repeated the analysis using the TIGRFAM database (Li et al., 2021), which contains a dedicated HMM profile for PurZ. Consistent with the Pfam-based analysis, the PurZ TIGRFAM domain ranked highest among all protein domains (**Supplementary Fig. S2A**). This result indicates that the enrichment strategy, in combination with the metaGPA pipeline, effectively captures dZ phages and identifies protein families central to the dZ biosynthetic pathway.

The distinction between *purA* and *purZ* is also reflected by the evolutionary relationships among genes containing the adenylosuccinate synthetase domain from both our dataset and the literature. Indeed, a phylogenetic analysis on a total of 46 complete CDSs containing a Pfam annotated adenylosuccinate synthetase domain revealed that most of the proteins from the dZ modified contigs (28 out of 29) clustered with the known PurZs while those from predicted canonical contigs (14 out of 17) formed a distinct clade together with the *E. coli* PurA (**Fig. 3A**). Taken together, these results are consistent with the predicted differential substrate specificity among these proteins and further support that our selection using REase treatment differentiate PurZ from the rest of adenylosuccinate synthetases superfamily.

**Figure 3.**
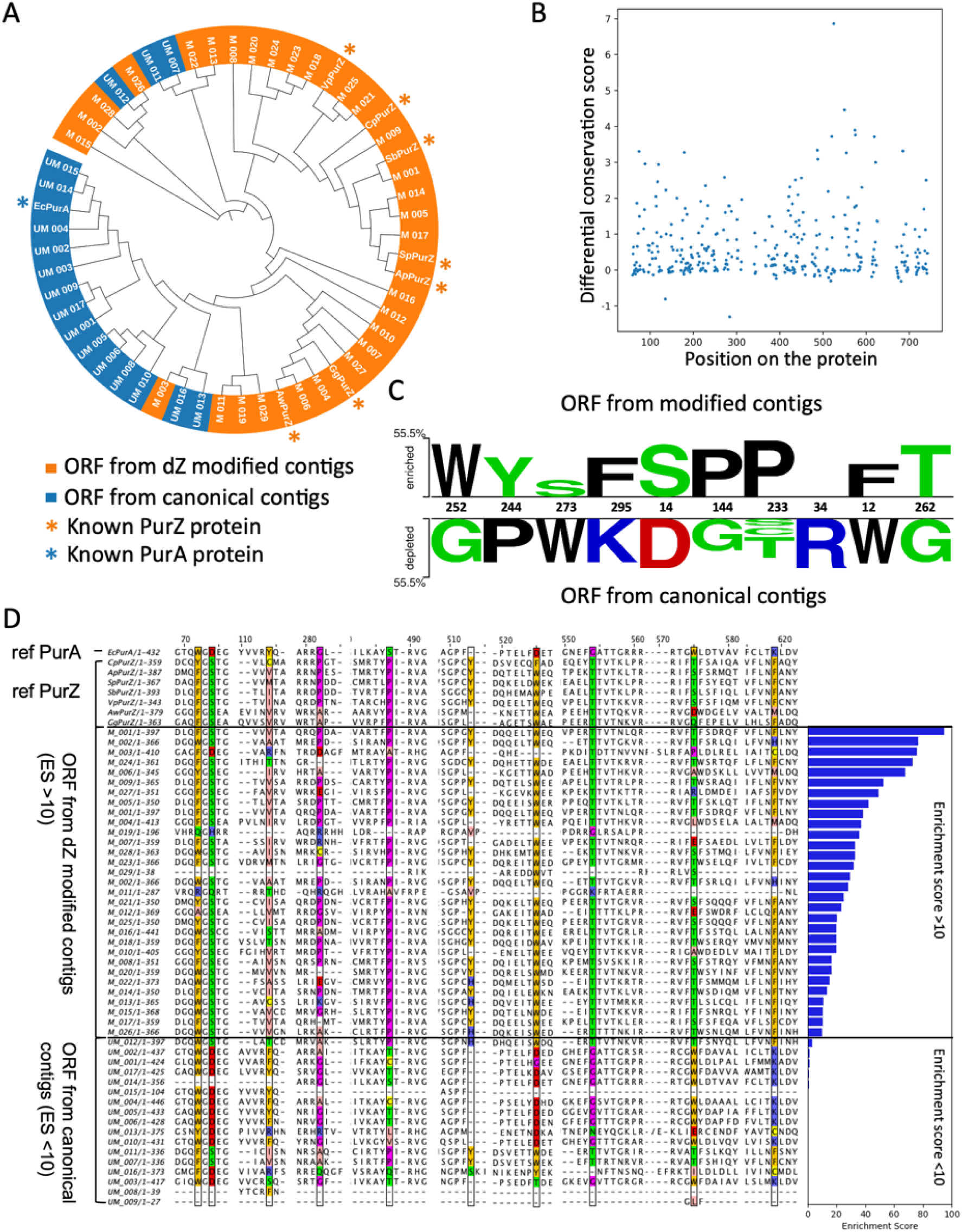
Phylogeny of PurZ and prediction of key amino acid residues. **A**) Phylogenetic tree of genes containing an adenylosuccinate synthetase domain from this study along with known *purA* and *purZ* genes. ORF from dZ modified contigs were colored in orange and ORF from canonical contigs were colored in blue. A few known PurZ and PurA reference “markers” were included in the alignment to provide evolutionary context. These markers contain the *E. coli* PurA and seven previously characterized phage PurZs, including *Cyanophage S-2L* PurZ (*Cp*PurZ), *Acinetobacgter* phage SH-Ab 15497 PurZ (*Ap*PurZ), *Salmonella* phage PMBT28 PurZ (*Sp*PurZ), *Sinbacteraceae* bacterium phage PurZ (*Sb*PurZ), *Vibrio* phage PhiVC8 PurZ (*Vp*PurZ), *Arthrobacter* phage Wayne PurZ (*Aw*PurZ) and *Gordonia* phage Ghobes PurZ (*Gg*PurZ). **B**) Differential conservation score of residues in predicted PurZ from modified contigs. **C**) Sequence logo of the top 10 residues with the most significant differential conservation score. Top and bottom panel residues are from predicted ORF from the modified and canonical/unmodified contigs respectively. Residue numberings are relative to PurZ in *Vibro* phage PhiVC8 (*Vp*PurZ). The two sample logo (14) program with default parameters was used to generate the figure (residues are shown if p-value < 0.05). **D**) Subset of the multiple sequence alignments of adenylosuccinate synthetases highlighting the top scoring residues colored by their physicochemical properties using zappo color scheme. The PurZ and PurA reference markers above were added to the alignments for comparison but not used in the calculation of the differential conservation score.

**Supplementary Figure S1.**
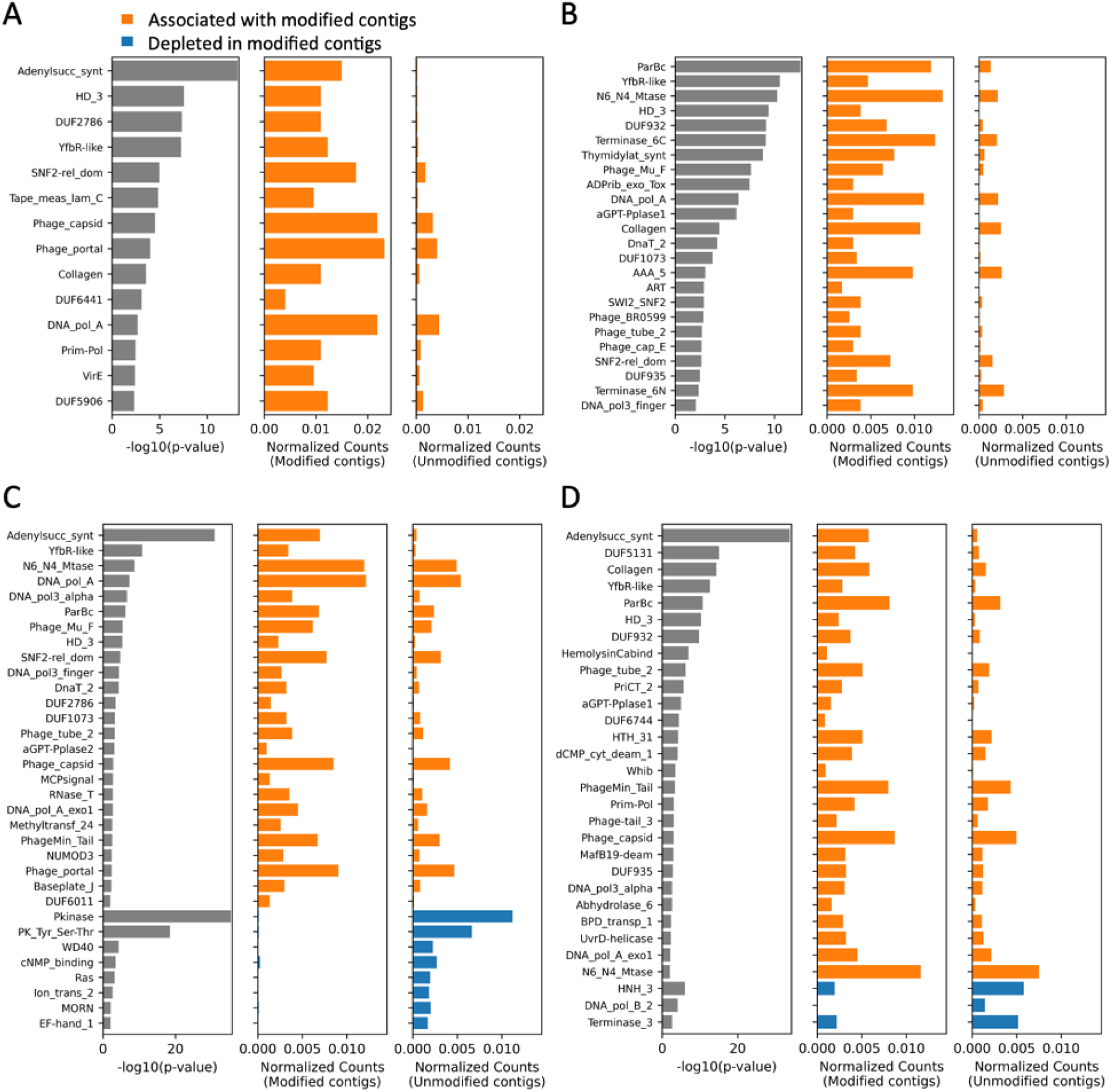
metaGPA identifies Pfams associated with the dZ modification. Pfams significantly (two-tailed p value < 0.01) associated with dZ modification in (**A**) ocean, (**B**) river and (**C-D**) two salt-marsh microbiome samples. Left: p value of each Pfam association analysis; middle: normalized counts in metaGPA predicted dZ-modified contigs; right: normalized counts in unmodified contigs. Enriched Pfams in modified contigs are marked in orange and depleted Pfams in modified contigs are marked in blue.

**Supplementary Figure S2.**
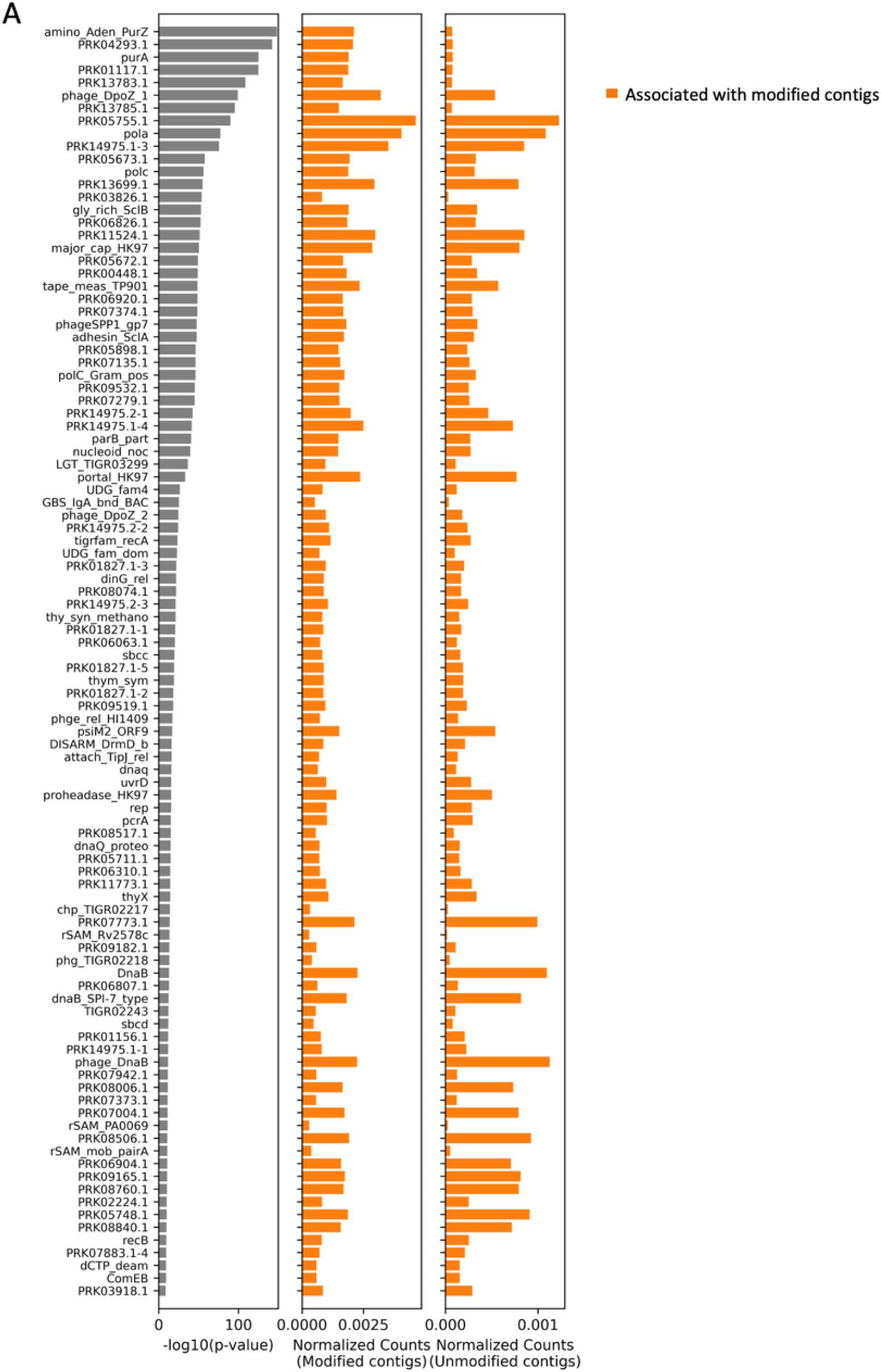
metaGPA identifies TIGRFAMs associated with the dZ modification. **A**) Top 100 TIGRFAM entries associated with the dZ modification in the composite dataset. Left: two-tailed p value of each TIGRFAM association analysis; middle: normalized counts in metaGPA predicted dZ modified contigs; right: normalized counts in unmodified contigs. Enriched Pfams in modified contigs are marked in orange and depleted Pfams in modified contigs are marked in blue.

### 4. metaGPA predicts key residues in the PurZ family involved in dZ formation

We have previously shown that the metaGPA framework can identify key residues involved in substrate specificity through differential residue conservation analysis. (11). Here, we applied the differential residue conservation analysis to the adenylosuccinate synthetase domain to further investigate potential key residues distinguishing dZ biosynthesis activity (PurZ) from PurA. We aligned the 46 protein sequences containing a predicted adenylosuccinate synthetase domain, grouping them into protein sequences from dZ-modified contigs (29 predicted PurZ proteins) or protein sequences from unmodified dA contigs (17 predicted PurA proteins). The top 10 residues exhibiting differential conservation signatures between the two groups were identified and selected for further investigation (**Fig. 3B-D, Supplementary Table 1**). Among these, **F295** and **Y298** (*Vp*PurZ coordinates) have been previously identified to be within the NTP catalytic binding site (3, 5) (**Fig. 3C-D, Supplementary Table 1**). The presence of these two residues has been shown to be important for stabilizing the adenine ring of ATP (3, 5). Additionally, **S14, F12,** and **T262** have been shown to accommodate the 2-amino group of dGMP (**Fig. 3C-D**, **Supplementary Table 1**). Notably, mutation of **S14 to D14**, the corresponding residue in *E. coli* PurA, has been shown to abolish substrate selectivity (5).

Besides the residues with characterized function, an unstructured region facing the catalytic pocket containing **G256** was reported by earlier work (3). The same group also showed that **P233** was located near the catalytic site which may suggest substrate selectivity. Our metaGPA analysis corroborates previously identified functionally validated amino acid residues and identifies additional novel residue candidates that may contribute to enzyme selectivity, including **W252** (ranking first), **Y244** (ranking second), **S273** (ranking third), **P144** (ranking sixth) and **I34** (ranking eighth) (**Supplementary Table 1**), though further experiments are needed to validate their potential activity.

**Supplementary Table 1.** Top 20 differentially conserved amino acid residues and corresponding aa residues in *Vp*PurZ and *Ec*PurA. The 10 entries highlighted in bold were chosen for further analysis (see Fig. 2).

| position | fraction_present | differential_<br>conservation_score | VpPurZ<br>position | VpPurZ<br>residue | EcPurA<br>position | EcPurA<br>residue |
| --- | --- | --- | --- | --- | --- | --- |
| 525 | 0.90740741 | 2.8655979 | **252** | **W** | **288** | **G** |
| 513 | 0.7962963 | 2.71853547 | **244** | **Y** | **280** | **P** |
| 575 | 0.90740741 | 2.38427816 | **273** | **S** | **310** | **W** |
| 619 | 0.88888889 | 1.92669869 | **295** | **F** | **332** | **K** |
| 74 | 0.90740741 | 1.89163858 | **14** | **S** | **14** | **D** |
| 282 | 0.94444444 | 1.86273345 | **144** | **P** | **146** | **G** |
| 487 | 0.92592593 | 1.80548699 | **233** | **P** | **271** | **S** |
| 114 | 0.92592593 | 1.79573071 | **34** | **I** | **33** | **R** |
| 72 | 0.90740741 | 1.76708582 | **12** | **F** | **12** | **W** |
| 554 | 0.92592593 | 1.73829099 | **262** | **T** | **299** | **G** |
| 179 | 0.88888889 | 1.72340574 | 74 | A | 71 | N |
| 622 | 0.88888889 | 1.71819479 | 298 | Y | 335 | V |
| 85 | 0.90740741 | 1.71722842 | 19 | L | 19 | K |
| 450 | 0.90740741 | 1.67743764 | 214 | D | 252 | G |
| 681 | 0.57407407 | 1.66011905 | - | - | 393 | P |
| 542 | 0.90740741 | 1.56774695 | 256 | G | 292 | C |
| 574 | 0.88888889 | 1.54132554 | 272 | F | 309 | G |
| 445 | 0.90740741 | 1.53677249 | 213 | A | 251 | T |
| 421 | 0.90740741 | 1.50079365 | 190 | S | 228 | L |
| 442 | 0.90740741 | 1.49758125 | 210 | R | 248 | G |

### 5. metaGPA co-occurrence network analysis recapitulates the entire phage-encoded dZ biosynthetic pathway

In addition to the adenylosuccinate synthetase domain, a total of 116 protein domain families were identified as significantly associated with dZ-modified DNA (p-value <0.01, **Fig. 2A**). To further examine their relationship, we performed a network analysis based on co-occurence of these domains on dZ-modified contigs (**Fig. 4A**). Six domains were directly connected with the adenylosuccinate synthetase domain and among those, five were also found in close genomic proximity (within 3 kb) to a predicted purZ gene (**Fig. 4B**). These include the HD_3 domain, the dATP/dGTP diphosphohydrolase N-terminal domain, and the DNA polymerase A/exo1 domains which are all known to be key components of the dZ biosynthetic pathway (2, 4, 5) The dATP/dGTP pyrophosphohydrolase domain (Pfam accession: PF04447) present in the MazZ enzyme has been shown to catalyze the hydrolysis of dATP/dGTP to dAMP/dGMP; the YfbR-like and HD_3 domains (Pfam accession: PF12917 and PF13023, both domains are overlapping) present in dATPase/datZ have been shown to remove dATP and dADP from the nucleotide pool of the host; and the DNA polA and DNA polA 3’-5’ exonuclease domains (Pfam accessions: PF00476 and PF01612) present in DNA polymerases have been shown to preferentially select for dZTP instead of dATP during replication; and Wayne-like DpoZ clade genomes seem to encode a gene homologous to dUTPase (Pfam accession: PF00692) to fulfill the role of DatZ and MazZ (8) in support of the dZ biosynthesis.

**Figure 4.**
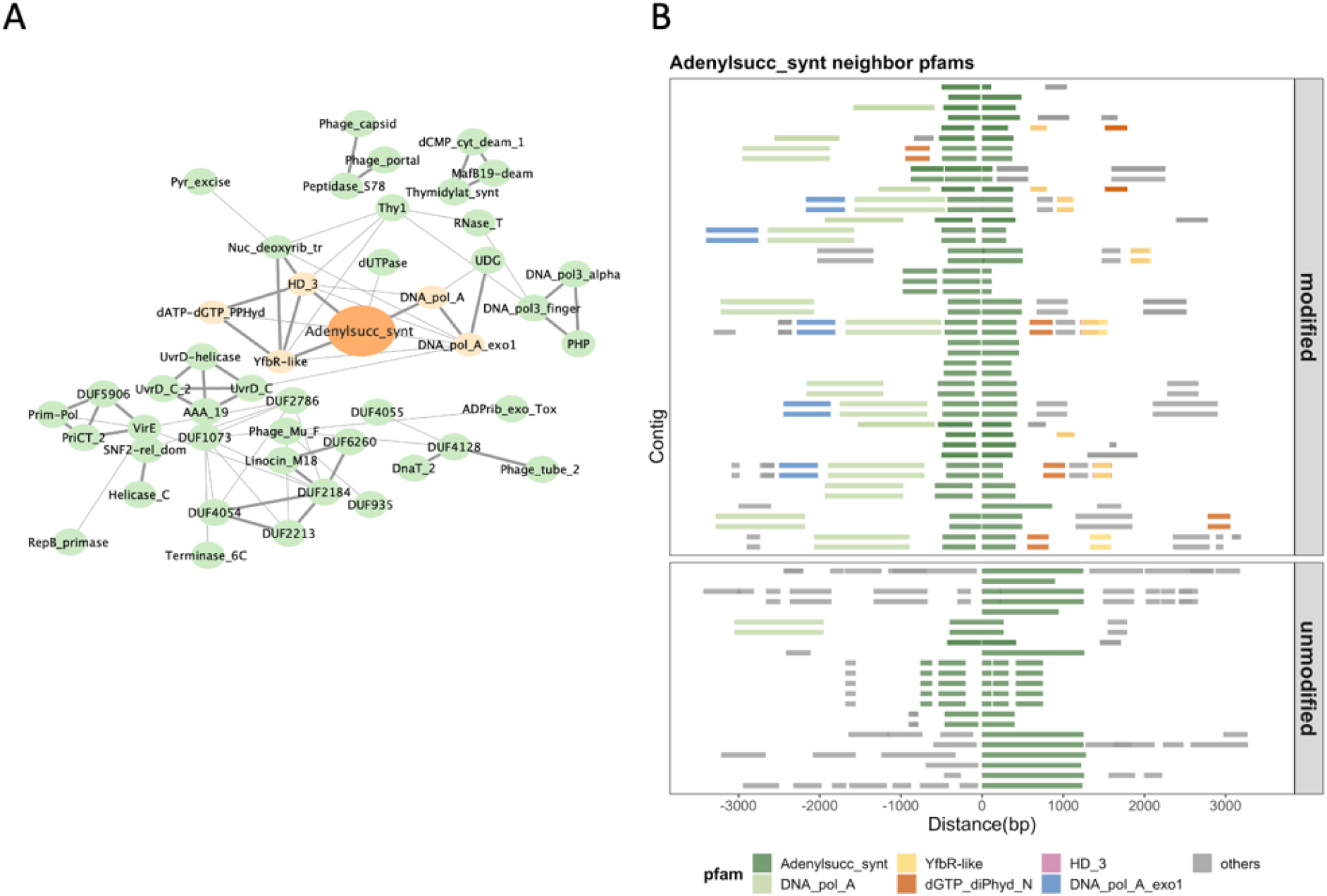
metaGPA recapitulates the dZ biosynthetic pathway. **A**) Co-occurrence networks of the 116 top associated Pfams (p-value < 0.01) that are directly or indirectly linked to the adenosuccinate synthase domain. Length and thickness of the edges reflect the significance of the associations, with thicker and shorter edges indicating stronger association. Only positive correlations with co-occurrence p-values < 0.01 are displayed. Key interactions connected to the adenylosuccinate synthetase domain are highlighted in orange. **B**) Genomic context in proximity to adenylosuccinate synthetases. Adenylosuccinate synthetase is centered in the middle and neighboring Pfams spanning 3 kb upstream and downstream (+/- 3 kb) are displayed as solid squares. Colored blocks show the top 5 Pfams/genes co-localized with annotated adenylosuccinate synthetases in metaGPA predicted dZ-modified contig.

Both the network analysis and genomic proximity are consistent with previous reports, highlighting the entire phage-encoded enzymes/domains and their roles in the dZ biosynthesis pathway.

### 6. metaGPA suggest alternative residues involved in substrate specificity in DatZ

Previous work by (2) has shown that substrate specificity in DatZ is primarily determined by two key residues: isoleucine at position 22 (**I22**), which provides specificity towards the adenine nucleobase, and tryptophan at position 20 (**W20**), which confers specificity for the sugar moiety. Building on these findings, we analyzed the corresponding positions across all annotated proteins containing the YfbR-like domain that define DatZ. Consistent with the findings of Czernecki et al., the **W20**-**I22** combination was observed exclusively in the YfbR-like domain found in the dZ-modified contigs with an enrichment score above 10 (19 out of 36, **Supplementary Fig. S3**). Interestingly, alternative residues at position 22, such as threonine (T) or serine (S), were also prevalent in annotated enzymes from enriched contigs, often in combination with tyrosine at position 20 (**Y20**). This Y20-T/S22 combination may suggest an alternative mechanism by which the protein achieves sugar-nucleobase specificity.

**Supplementary Figure S3.**
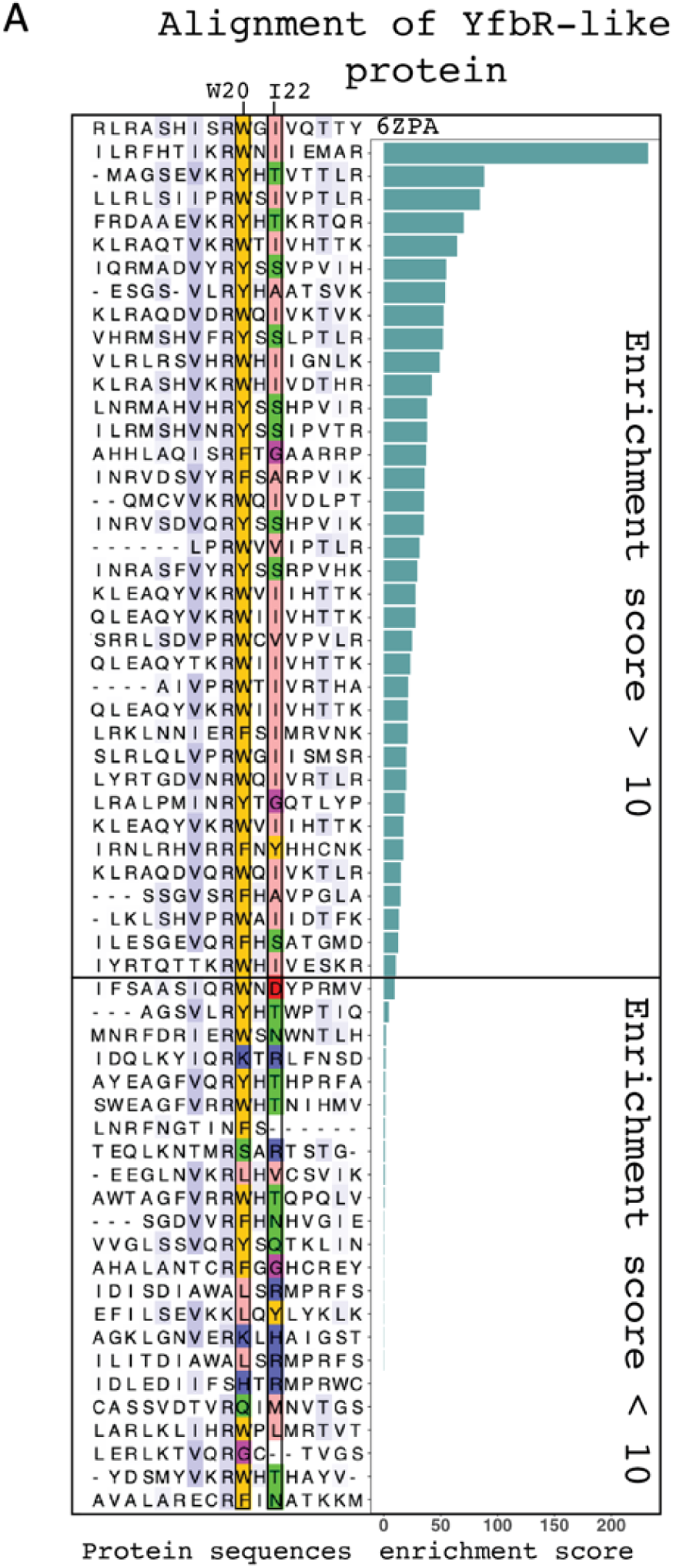
Alignment of the YfbR-like domain. Section of an alignment of the Cyanophage S-2L HD phosphohydrolase DatZ with the predicted proteins to contain the YfbR-like domain. The two key residues in DatZ **W20** and **I22** (PDB: 6ZPA) are highlighted. Sequences were ranked according to the enrichment score in metaGPA.

### 7. Validation of NDT as an novel potential component of the dZ biosynthetic pathway with preferred substrate specificity for dA

The co-occurrence network of associated Pfam domains in dZ-modified contigs identifies two additional domains of interest: the nucleoside 2-deoxyribosyltransferase (NDT, Pfam accession: PF05014) and the uracil-DNA glycosylase (UDG, Pfam accession: PF03167). NDT which does not co-localize directly with adenylosuccinate synthetase domain but shows strong co-occurrence with DatZ (YfbR-like and HD_3 domains) and DNA pol A, suggesting a yet unknown link to the dZ biosynthetic pathway (**Fig. 2**, **Fig. 4B** and **Supplementary Fig. S2**). It has been reported that diaminopurine may affect glycosylase UDG binding by stabilizing DNA spontaneous opening with three hydrogen bonds (15). Taken together, these associations point toward a potential but yet undefined connection between the NDT and/or the UDG-containing proteins and the dZ biosynthetic pathway.

Given that the known function of NDT is to catalyze the transfer of the 2’ deoxyribosyl moiety from a donor deoxyribonucleoside to an acceptor purine or pyrimidine base (16), we hypothesized that the NDT enzymes found in dZ-modified phage may catalyze the exchange of purine bases between adenine and diaminopurine to further deplete dA in the nucleoside pool.

Further lines of evidence support this hypothesis. Phylogenetic analysis suggested clustering of NDT sequences from dZ modified contigs (**Supplementary Fig. S4A**) indicating a distinct function or substrate preferences compared to canonical NDT. To test this, we cloned all 10 NDT genes identified in dZ-modified contigs into T7 expression vectors. Two NDTs from canonical contigs (UM1 and UM6) as well as a previously characterized control NDT from *Lactobacillus delbrueckii* (*Ld*NDT) were also included (**Supplementary Fig. S4B**) (17–19). Five of the ten novel NDTs expressed as soluble protein and two (M2 and M3) were randomly picked for purification (**Supplementary Fig. S4B** and **Supplementary Fig. S5A-B**). Transglycosylation reactions were carried out with purified NDTs using different pairs of 2-deoxyribonucleoside donors and nucleobase acceptors. As expected, the control *Ld*NDT exhibited activity over a broad range of donor/acceptor combinations (**Fig. 5A-D, Supplementary Fig. S6A-E and Supplementary Table 2**). NDT-M2 reacted with both dA and dZ substrates as 2-deoxyribonucleoside donors (**Fig. 5A-B, Supplementary Table 2**). The preference for dA was on average 3.4-fold higher than for dZ (**Fig. 5E, Supplementary Table 2**). NDT-M2 also showed activity with dG, however, a 37-fold higher preference for dA over dG as the 2-deoxyribonucleoside donor was seen (**Fig. 5C-E, Supplementary Table 2**). NDT-M3 and the other two NDTs from predicted unmodified genomes didn’t show transglycosylation reactions with dA, dG or dZ (**Fig. 5A-D**). We also tested the reactions of NDT-M2 with pyrimidine 2-deoxyribonucleoside donor and purine nucleobase acceptors but observed no activity (**Supplementary Fig. S6C-E, Supplementary Table 2**). Together, the results suggest that NDT-M2 functions as a type I purine/purine specific 2-deoxyribosyltransferase with preferred dA activity. This substrate preference supports the proposed role of NDT-M2 in depleting the dA pool in favor of the dZ biosynthesis.

**Figure 5.**
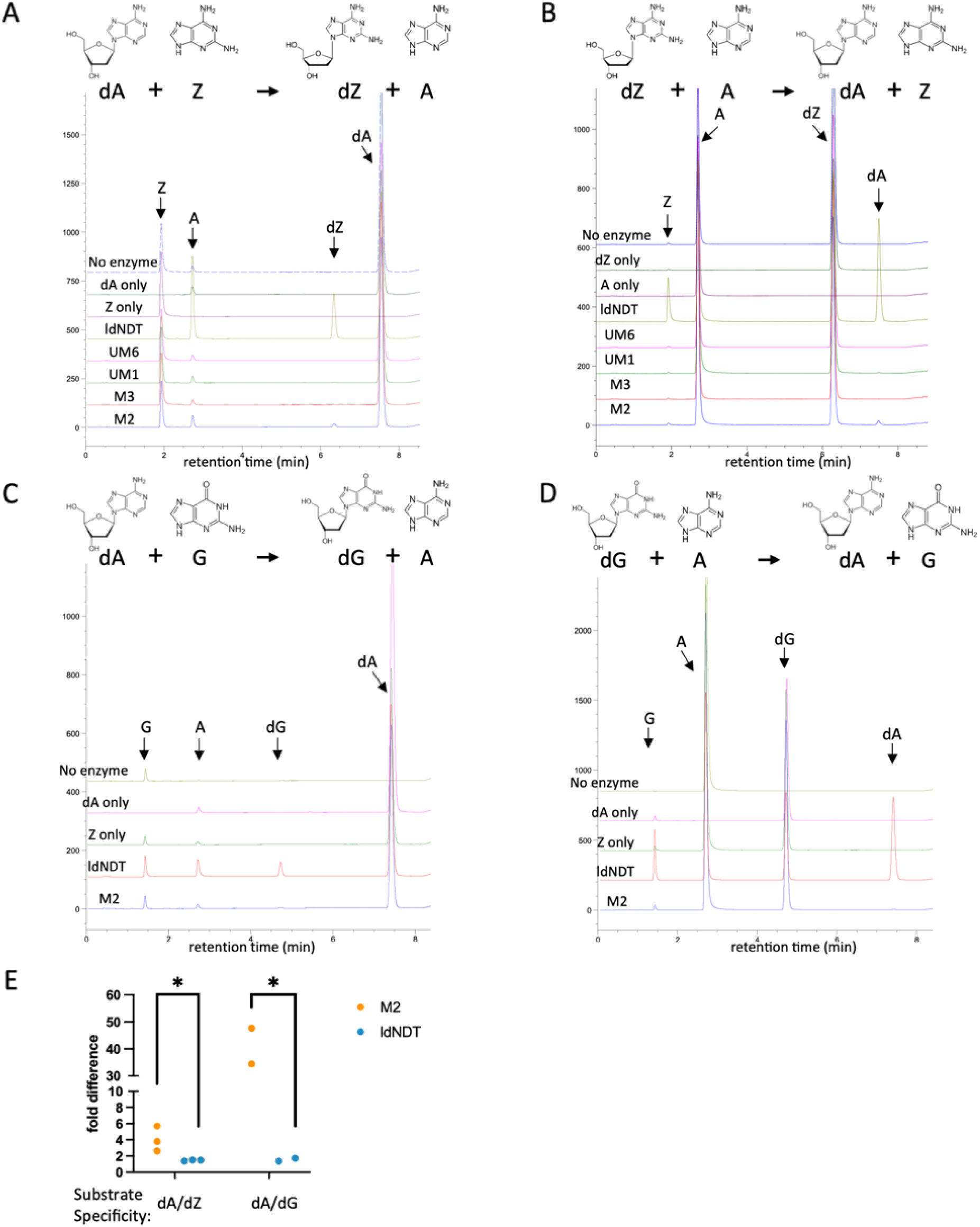
Validation of NDT activity towards dA over dZ. **A**) HPLC profile illustrating the enzymatic activity of NDT in presence of dA and Z. For each reaction, an equal molar concentration of nucleoside donor and nucleobase acceptor were added. dA only: denote the reaction containing only the donor dA, but no acceptor Z. *Ld*NDT enzyme was added into the reaction. Z only: reaction contained only the acceptor Z (no donor dA). No enzyme: reaction contained both donor and acceptor but no enzyme. UM1,UM6, M3 and M2 are the purified enzymes. **B**) Same as in **A)** but using dZ and A. **C**) Same as in **A)** but using dA and G. **D**) Same as in **A)** but using dG and A. **E**) Substrate preference of *ld*NDT (blue) and M2-NDT (orange). The ratio differences in enzyme activities using either dA over dZ or dA over dG as donors were measured with 2-3 biological experiment repeats. Asterisks denote p value < 0.05.

**Supplementary Figure 4.**
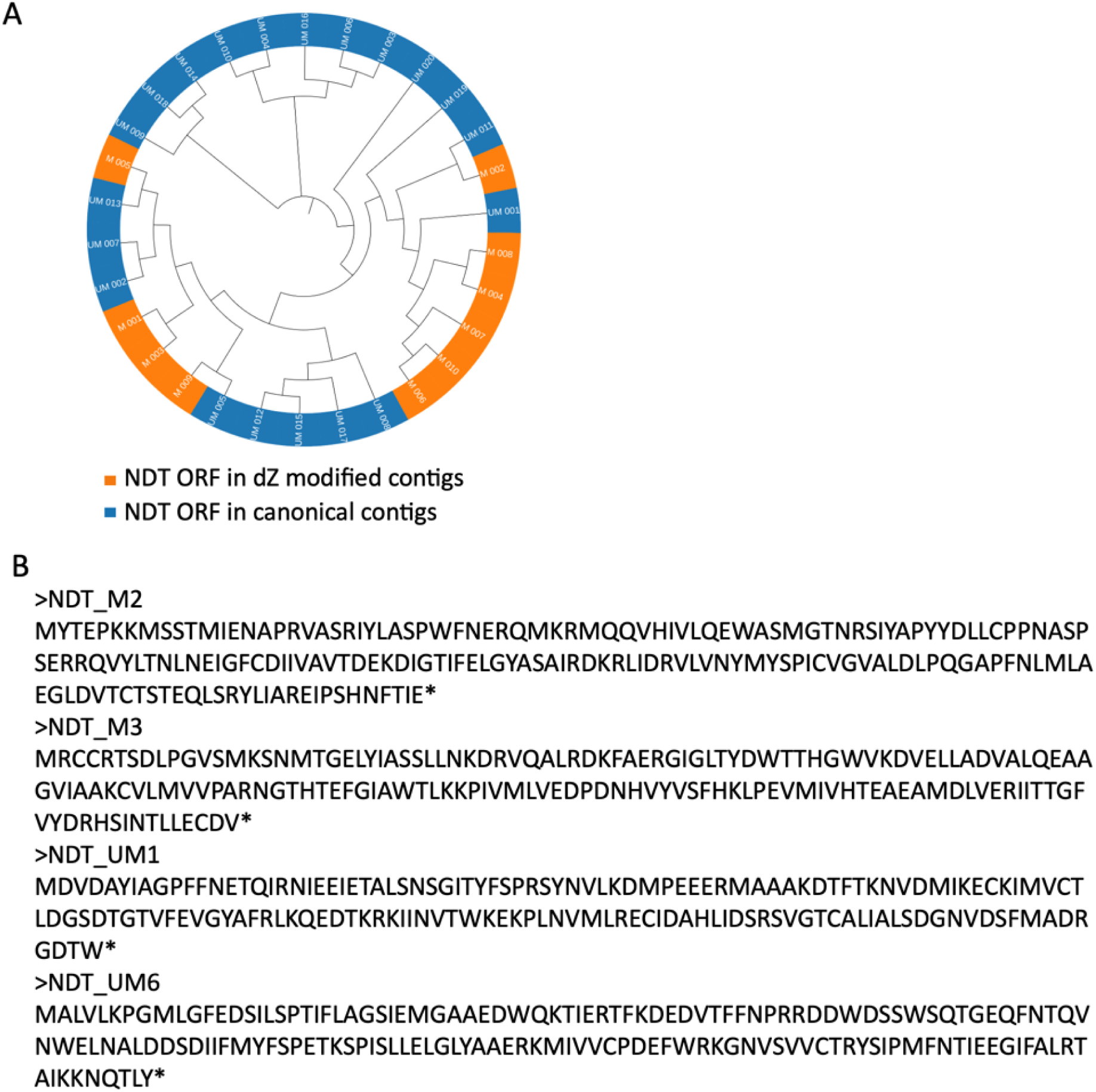
*de novo* NDT identification from metaGPA. **A**) Phylogenetic tree of NDT domains (Pfam accession: PF05014) from metaGPA analysis. Orange indicates NDT domain identified in metaGPA predicted dZ-modified contigs; Blue indicates NDT domain identified in canonical contigs. **B**) Protein sequences of used NDTs in this study.

**Supplementary Figure S5.**
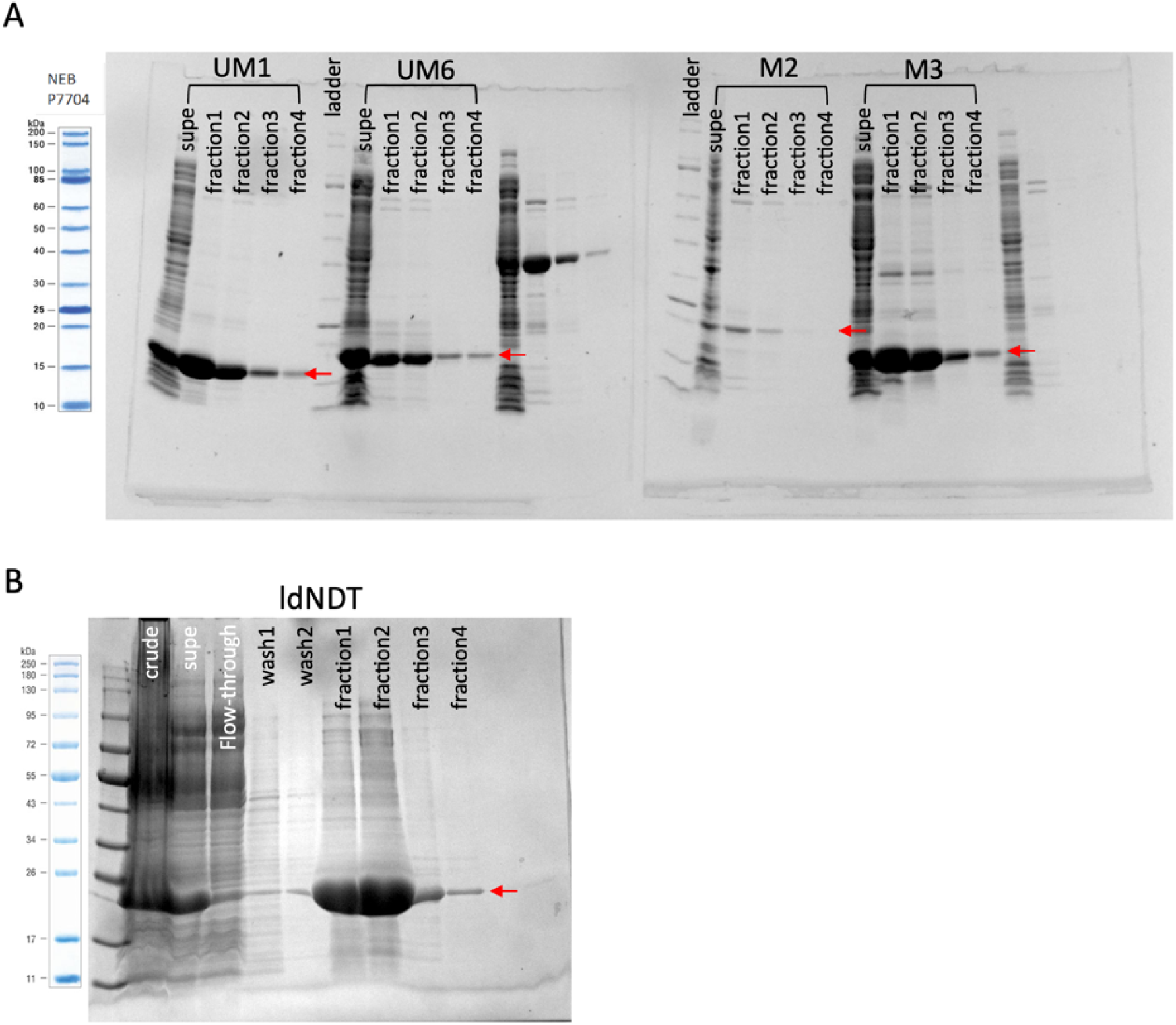
Expression and purification of NDTs. **A**) SDS-PAGE gel showing purification of the four NDT proteins identified in this study. Expected molecular weights of the proteins are indicated by red arrows. **B**) SDS-PAGE gel showing purification of control *Ld*NDT protein. Expected protein molecular weights are indicated by red arrows.

**Supplementary Figure S6.**
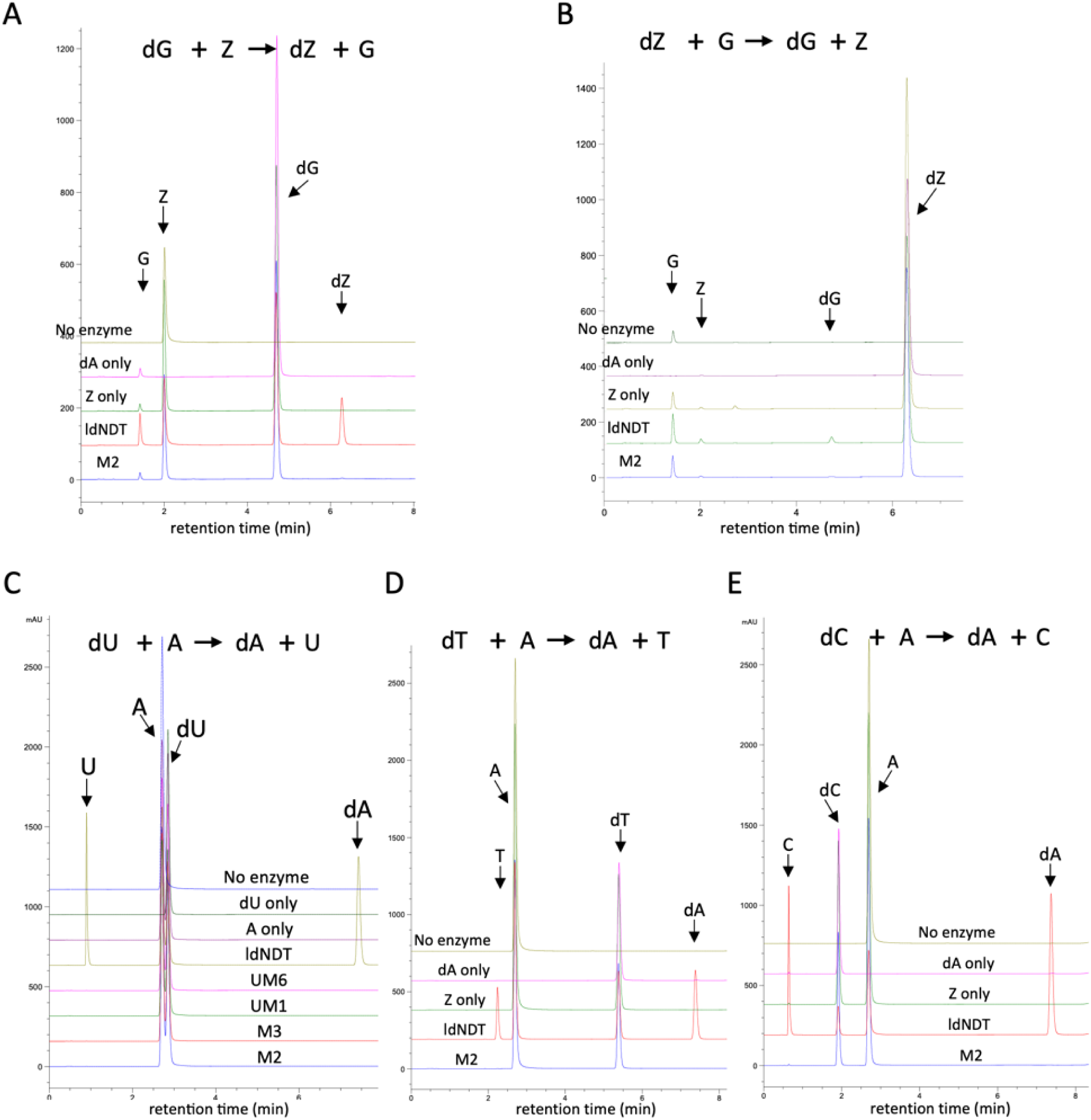
Validation of NDT activity. **A**) HPLC profile illustrating the enzymatic activity of NDT in presence of dG and Z. For each reaction, an equimolar concentration of nucleoside donor and nucleobase acceptor were added. dG only: reaction contained only the donor dG but no acceptor A. *Ld*NDT enzyme was added into the reaction. Z only: reaction contained only the acceptor Z (no donor dG). No enzyme: reaction contained both donor and acceptor but no enzyme. **B**) same as in **A)** but using dZ and G. **D**) same as in **A)** but using dT and A. **E**) same as in **A)** but using dC and A.

**Supplementary Table 2.** Quantification of NDT activity on different substrates.

| Reaction category | Nucleoside donor | Nucleobase acceptor | IdNDT average conversion | M2 average conversion |
| --- | --- | --- | --- | --- |
| Purine ↔ purine | dA | Z | 48.4% | 3.4% |
| Purine ↔ purine | dZ | A | 33.5% | 1.0% |
| Purine ↔ purine | dA | G | 58.1% | 11.2% |
| Purine ↔ purine | dG | A | 37.4% | 0.3% |
| Purine ↔ purine | dG | Z | 30.4% | 0.5% |
| Purine ↔ purine | dZ | G | 30.8% | 3.8% |
| Purine ↔ pyrimidine | dU | A | 47.3% | - |
| Purine ↔ pyrimidine | dC | A | 66.8% | - |
| Purine ↔ pyrimidine | dT | A | 33.9% | - |
| Purine ↔ pyrimidine | dC | Z | 72.9% | - |
| Purine ↔ pyrimidine | dT | Z | 35.3% | - |
| Purine ↔ pyrimidine | dA | T | 21.9% | - |
| Purine ↔ pyrimidine | dA | C | 27.6% | - |

### 8. Novel DNA polymerases identified from dZ modified microbiome promotes *in vivo* incorporation of dZ

It has been previously shown that dZ phages encode a noncanonical homolog of DNA polymerase A, DpoZ, that preferentially select for aminoadenine deoxynucleoside triphosphate (dZTP) instead of adenine deoxynucleoside triphosphate (dATP) during replication (4). In our study, we identified a few novel DpoZ homologs from dZ modified contigs. Phylogenetic trees constructed with these homologs demonstrated divergent clustering, suggesting their functional differentiation from the canonical DNA pol A (**Supplementary Fig. S7A**). We cloned two of these novel DpoZ homologs assembled from the environmental phage microbiome samples, namely phiDpoZ1 and phiDpoZ2 (**Supplementary Fig. S7B**). Both of them share structural conservation with *Vp*DpoZ (**Fig. 6A**). In a previous study, we demonstrated that *in vivo* dZ incorporation is possible by co-expression of *purZ*, *mazZ* and *datZ* (dZ cluster) in a plasmid. Introduction of *Sp*PurZ significantly increases the dZ incorporation level (9). Here we showed that the two novel phage DpoZ homologs, phiDpoZ1 and phiDpoZ2, when expressed together with the dZ cluster, also significantly elevated dZ incorporation, reaching levels comparable to those observed with *Sp*PurZ (**Fig. 6B**). We also cloned *Vp*PurZ and co-expressed it with the dZ cluster; however, no significant increase in dZ incorporation was observed (**Fig. 6B**). We then sought to further promote the level by co-expressing two more components: a helicase and a single strand DNA binding protein (SSB) that were cloned from the *Salmonella* phage PMBT28 genome (9). The addition of *Salmonella* phage PMBT28 helicase and SSB dramatically increased the dZ incorporation in recombinant plasmid to on average 53%. Combination of dZ cluster, phiDpoZ1 and *Salmonella* phage PMBT28 helicase-SSB slightly elevated the dZ incorporation to 34%, while phiDpoZ2 was not compatible with *Salmonella* phage PMBT28 helicase-SSB (**Fig. 6C**). It is possible that a matching helicase and SSB or other additional component from the same phage genome may be required for increased DpoZ activity. These results validated novel DNA pol A homolog DpoZs with preferential incorporation of the dZ base into the genome.

**Figure 6.**
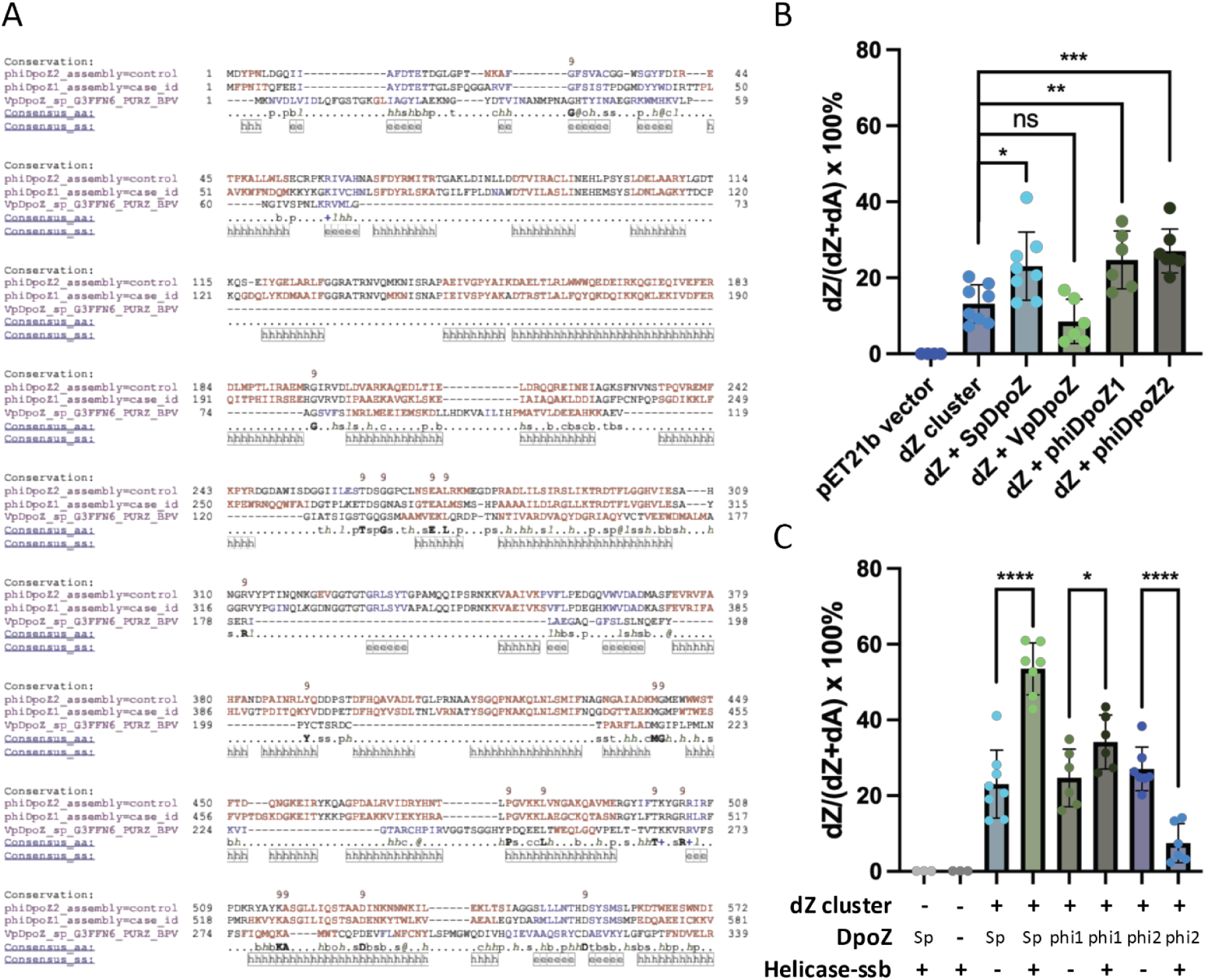
metaGPA-identified DpoZs increases dZ incorporation *in vivo* when co-expressed with dZ cluster. **A**) Structural multi-alignment between *Vp*DpoZ and *de novo* DpoZs (phiDpoZ1 and phiDpoZ2) from metaGPA study. Predicted secondary structures are colored (red: alpha-helix; blue: beta-strand). Multiple sequence alignment and structural prediction are generated with PROMALS3D (17). **B**) dZ incorporation levels in *E.coli* from co-expression of *Salmonella* phage PMBT28 dZ cluster (*purZ*, *mazZ* and *datZ)* and DpoZ homologs. Data from two biological repeats (n=6-8) are shown. ns: p value not significant; *: p-value < 0.05; **: p-value < 0.01; ***: p-value < 0.001; ****: p-value < 0.0001. **C**) dZ incorporation levels from co-expression of *Salmonella* phage PMBT28 dZ cluster (*purZ*, *mazZ* and *datZ)*, *Salmonella* phage PMBT28 helicase and SSB, DpoZ homologs. Data from two biological repeats (n=6-8) are shown. *: p-value < 0.05; ****: p-value < 0.0001.

**Supplementary Figure S7.**
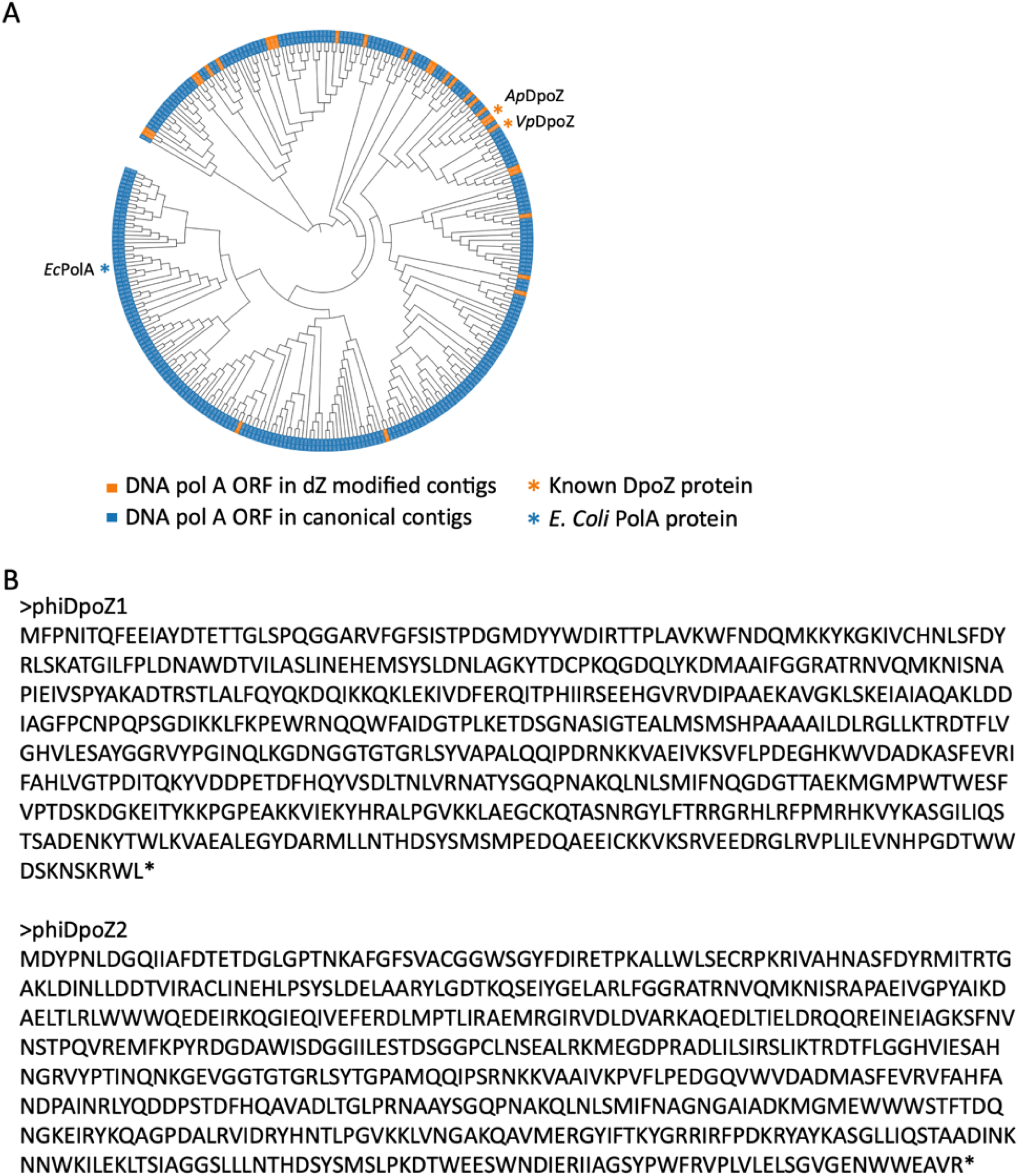
metaGPA-identified DpoZ homologs. **A**) Phylogenetic tree of DNA pol A domains (Pfam accession: PF00476) from metaGPA analysis. Orange indicates DNA pol A domains identified in metaGPA predicted dZ modified contigs; Blue indicates DNA pol A domain identified in canonical contigs. Known DpoZ and canonical PolA reference “markers” were included in the alignment to provide evolutionary context. These markers contain the *E. coli* Klenow PolA and two previously characterized phage DpoZs, including *Acinetobacgter* phage SH-Ab 15497 DpoZ (*Ap*PurZ) and *Vibrio* phage PhiVC8 DpoZ (*Vp*DpoZ). **B**) Protein sequence of the two novel DpoZ homologs validated in this study.

## Discussion

In this paper, we present the first study shown to effectively select dZ-containing genomic DNA directly from the viral-fraction of aquatic microbiomes. This method processes the bulk genomic DNA, bypassing the conventional and laborious process of isolating and culturing individual phage strains. Utilizing the resistance of the dZ DNA modification from digestion by a designated mixture of restriction enzymes, gDNA from phages without the dZ modification can be greatly diminished, enhancing sequencing throughput utilization for positive dZ phage genome signals in a cost-effective manner. Moreover, this strategy can be extended to other applications including other DNA nucleobase modifications or other microbiome organisms. The idea of exploiting methylation-sensitive restriction enzymes to digest unmethylated DNA and enriching certain bacterial taxa of interest from microbiome has been tested and validated by others (mEnrich-seq and REMoDE) (20, 21), demonstrating the potential and flexibility of this strategy to explore the diversity of microbiome. One caveat is that the selection may lead to any modified DNA (e.g. 6mA modified DNA) that is partially or fully resistant to the REase cocktail used in the enrichment/selection step. Possible solutions to improve the specificity of selection could include applying more stringent selection reagents or setting up more stringent analysis cutoffs (e.g. increasing the enrichment score).

Linking the phenotype identified through selection to the corresponding genotype using metaGPA provides an effective approach to uncover biosynthetic pathways associated with the phenotype (11). In this study, we recapitulated the major enzymatic components in the biosynthesis of dZ in one single experiment. In contrast, conventional approaches to gene and pathway biochemical characterization often involve substantially more labor-intensive and resource-demanding processes.

Our study reveals discoveries at two distinct levels: First, for protein families with previously characterized roles in the dZ pathways, metaGPA stratifies broad protein domains into functionally refined categories. metaGPA allows for the delineation of key amino acid residues unique to PurZ and those conserved in canonical PurA. Consequently, proteins containing the adenylosuccinate synthase domain can be functionally subdivided into the canonical PurA for the synthesis of AMP and PurZ for the synthesis of ZMP. In contrast, the Pfam annotation of the adenylosuccinate synthase domains alone is insufficient to distinguish between these two functional classes. We have also specified and experimentally validated a subgroup of DNA polymerases that enhances dZ incorporation, thus contributing to the biosynthesis and maintenance of the dZ genomes. Finally, we discovered that the Y20-T22/S22 combination in the YfbR-like domain is uniquely observed in predicted dZ contigs. The potential role of this motif in determining nucleotide hydrolysis specificity, however, remains to be investigated.

The second level of discoveries by metaGPA regards new components within a biological pathway. In this work, we identified NDT as a potential contributor in the dZ biosynthesis. NDT promotes selective nucleobase swap, thus effectively removing the dA nucleoside pool. As the assembly of genomes through metaGPA selection can be incomplete, we sought to search the ncbi virus database for more evidence of presence of NDT in dZ phage genomes (defined as containing PurZ homolog with S14 referred to vibriophage phiVC8 PurZ) (**Supplementary Fig. S8**). Interestingly, these dZ genomes all share genomic context features resembling Wayne-like dZ phage clade: they all lack the DatZ or MazZ in the genome, a gene homologous to dUTPase, instead, may fulfill the analogous role in dZ synthesis (4, 8). In addition, these genomes all contain the DpoZs (DNA primase-polymerase) which is distinct from the DNA polymerase I homologous DpoZs cyanophage S-2L or the Vibrio phage PhiVC8-like clade (4, 8). Given the NDT preference for dA, it is likely that in Wayne-like or homologous dZ genomes, NDT may be involved as an additional factor complementing dUTPase and DNA primase-polymerase DpoZs for the adenine exclusion in such genomes. Experimental validation to confirm the role of NDT in facilitating the specificity towards making 2-aminopurine genomes *in vivo* will need to be further verified. Elucidating the NDT-dUTPase involvement in alternative dZ synthesis could advance our understanding of the biochemical diversity in nucleic acid pathways and facilitate engineering efforts.

**Supplementary Figure S8.**
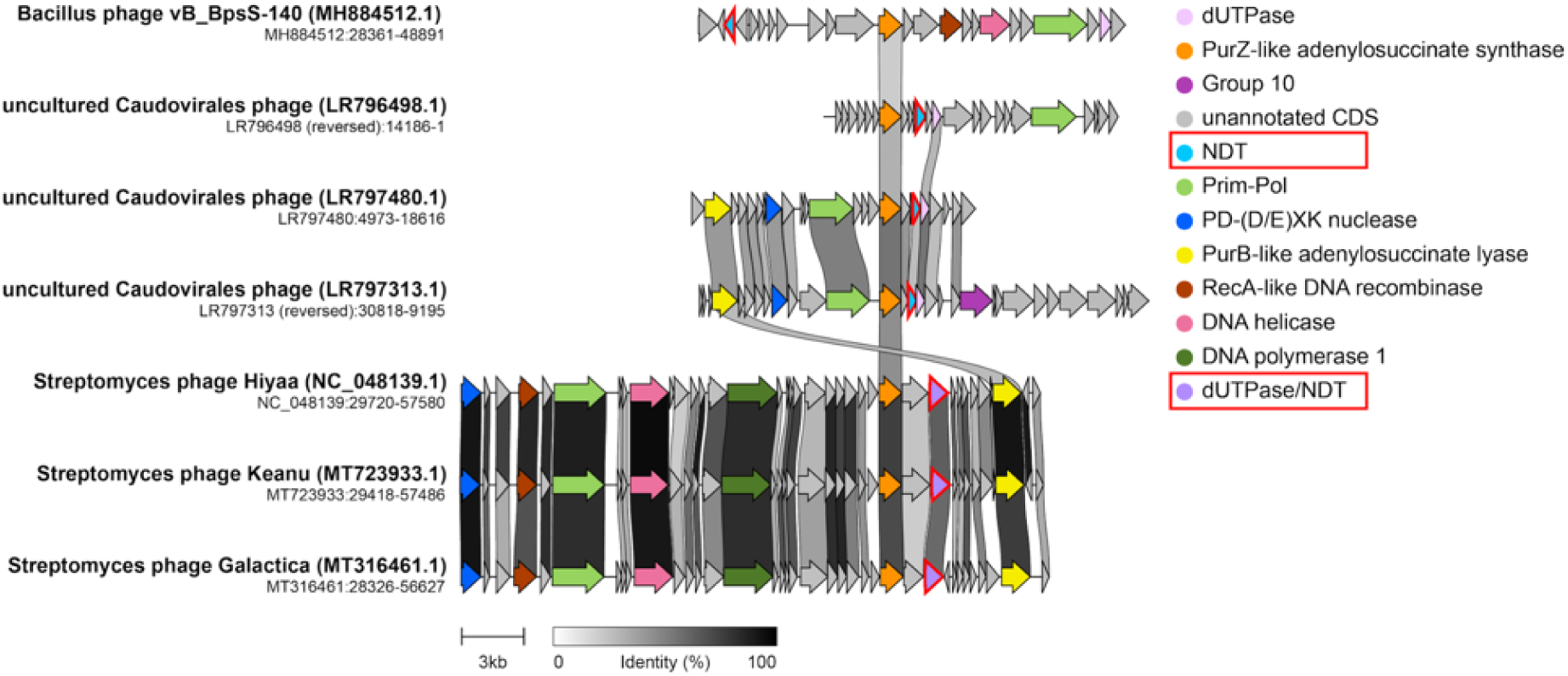
Co-localization of PurZ and NDT in genomes. The ORFs annotated as NDT or dUPTase/NDT dual protein are highlighted in red.

## Materials and Methods

### Metagenome sample collection and genomic DNA (gDNA) extraction

Four aquatic metagenome samples were collected at the coastal shore algae section near Beverly, MA; Ipswich river at Ipswich, MA and twice at the salt marsh at Hamlin reservation, MA respectively. For each sample, 2-4 L were used for phage collection: Large debris and bacterial cells were pelleted and removed using centrifugation at 5000 x g for 30 min at 4 °C. Phage particles in the supernatant were precipitated with 10% PEG8000 and 1 M NaCl, and resuspended in 2-4 mL of phage dilution buffer (10 mL Tris-HCl at pH 8.0, 10 mM MgCl2, 75 mM NaCl). We used phenol-chloroform DNA extraction protocol for genomic DNA extraction. Details of the protocol were described in our earlier publication (11).

#### Enzyme mix for dZ DNA enrichment

The seven enzymes and their recognition motifs are provided in the table below. In each individual restriction sensitivity/resistance test, PCR DNA using dZTP to replace dATP is fully resistant to each of these REases. W: A, T; R: A, G; Y: C, T.

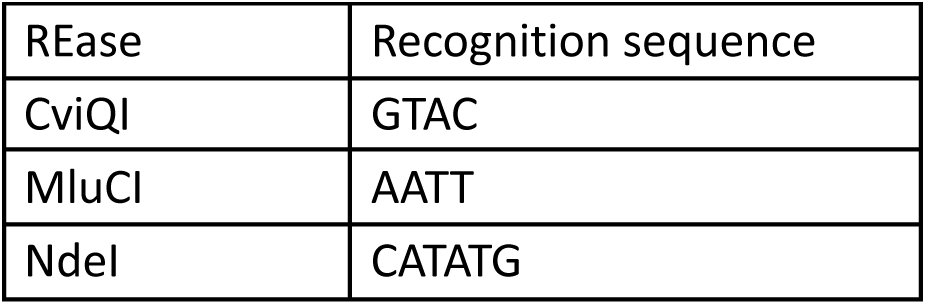
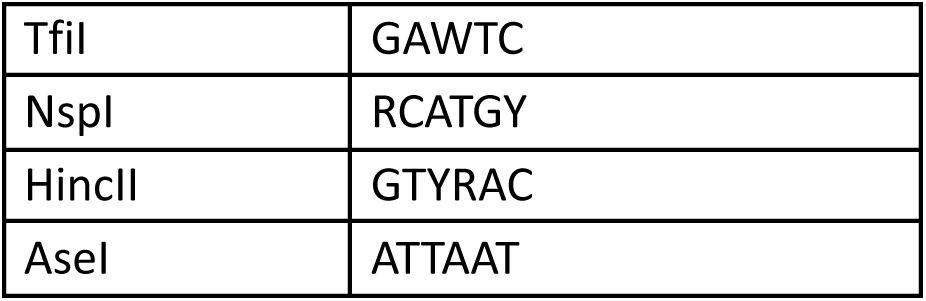

#### Enzymatic treatment to enrich dZ containing viral microbiome genomes

Extracted phage gDNA were divided into two groups: control and treatment. Each group contains 1.2 µg DNA input. For the enzyme treated group, DNA input was first treated with 0.5 µL NdeI (NEB cat: R0111S), 1 µL MluCI (NEB cat: R0538S), 1 µL NspI (NEB cat: R0602S), 1 µL HincII (NEB cat: R0103S), 1x cut smart buffer (NEB cat: B6004S) in 50 µL volume with 37°C incubation for 30 minutes. The products were heated to 65°C and incubated with 1 µL TfiI (NEB cat: R0546S) for 30 minutes. After that, the reaction was increased to 100 µL total volume in 1x NEBuffer 3 (NEB cat: B7003S) with 1 µL AseI (NEB cat: R0526S) and 1 µL CviQI (NEB cat:R0639S) added to the reaction. Incubation was conducted at sequential steps: 25°C for 15 minutes and then 37°C for 15 minutes. 5 µL Proteinase K (NEB cat: P8107S) were then added to inactivate the enzymes by heating to 37°C for 30 minutes. We used oligo clean & concentrator (Zymo, cat: D4060) to purify and elute digested DNA in 20 µL 0.1x TE buffer.

#### Size selection for dZ enriched DNA

Size selection was performed on both enzyme mix treated and control DNA. To prepare 35% v/v beads for size selection, 350 µL AMPure XP beads (Beckman Coulter, cat: A63880) were diluted with 650 µL 10 mM Tris pH 8.0. Enzyme mix treated or control DNA were then incubated with beads at 1 to 3.5 ratio by volume, with constant shaking for 15 minutes to facilitate binding. Beads and sample mixture were spun down briefly with a table centrifuge and placed in a magnetic rack. Unbound supernatant was removed using a pipette. The beads were washed twice with 80% ethanol and eluted in 50 µL 0.1x TE buffer for later DNA library preparation.

#### DNA library preparation and DNA sequencing

Prepared DNA samples were sheared to 200 bp in 50 μL 0.1x TE using Covaris S2 Focused Ultrasonicator. One reaction of NEBNext Ultra II DNA Library Prep Kit for Illumina (NEB cat: E7645) was used for each sample. The DNA library was purified with 1× volume of NEBNext Sample Purification Beads (NEB #E7103) and eluted with 40 μL of 1 mM Tris buffer (pH 7.5).

A second round enzyme treatment was performed on the treated group to further enrich dZ modified DNA. The products were purified with 1x volume AMPure beads and eluted in 40 µL 0.1x TE.

Finally, libraries were indexed (with NEBNext Multiplex Oligos for Illumina E6440), amplified using NEBNext Ultra II Q5 Master Mix (11 cycles for control library and 20 cycles for RE treated library) and pooled for sequencing on an Illumina Nextseq instrument with paired end reads of 75 bp.

#### qPCR for dZ enrichment efficiency

We spiked-in a 3 kb Z-DNA amplicon and E.coli MG1655 gDNA in the sheared phage DNA libraries to test REase mix digestion efficiency. After one round of REase mix treatment as described above, both treated and untreated control samples were purified using oligo clean & concentrator (Zymo, cat: D4060) and resuspended in 0.1X TE8. We then dilute the digestion product in water and performed qPCR with Luna Universal qPCR Master Mix (NEB #M3003). The following primers were used to quantify Z-DNA and E.coli DNA products. Results were normalized to untreated controls and a delta delta Ct method was adopted to quantify dZ enrichment efficiency.

E.coli-F: 5’-TTGCTGAGTTTCACGCTTGC

E.coli-R: 5’-AAAACCGCTTGTGGATTGCC

Z-DNA-loc1-F: 5’-CCTCGACCTGAATGGAAGCC

Z-DNA-loc1-R: 5’-GCGATGGATATGTTCTGCCAAGG

Z-DNA-loc2-R: 5’-CGAGCGGTATCAGCTCACTC

Z-DNA-loc2-R: 5’-CCACCTCTGACTTGAGCGTC

#### Data process and metaGPA analysis

Paired-end read FASTQ files were trimmed with Trim Galore v0.6.4 (https://www.bioinformatics.babraham.ac.uk/projects/trim_galore/) using the --paired option. MetaGPA code and analysis is available as described in (Yang et al. 2021). For this study, we used default parameters in MetaGPA main pipeline except for adjusting the contig enrichment score to 10 (-C 10). The Pfam database release 33.0 was used for annotation analysis.

#### Creation of the composite data set

We conducted four separate metaGPA analyses, one for each phage sample, using a consistent contig enrichment score threshold of 10 (-C 10). To construct a composite dataset, we concatenated all modified contigs (enrichment score ≥ 10) identified across the four individual datasets. Similarly, unmodified contigs (enrichment score < 10) from all samples were combined. Annotation of both modified and unmodified composite contigs was performed using the same pipeline. We then re-executed the Pfam domain association analysis (as implemented in the metaGPA pipeline) on the composite dataset, followed by phylogenetic analysis, residue differential conservation analysis and co-occurrence analyses, as described in the original metaGPA study (11).

#### Alignment and phylogenetic analysis

Full coding sequences of annotated protein domains identified from metaGPA pipeline or from reference database were aligned with multiple sequence alignment tool mafft v7.508 (Katoh and Standley 2013). The resulting aligned fasta files were subjected to construct phylogenetic trees using the maximum likelihood method in the phylogenetic analysis program RAxML v8.2.12 (Stamatakis 2014). We chose the -f a option to do rapid bootstrap analysis and the -m PROTGAMMAAUTO model to automatically determine the best protein substitution model to be used for the dataset. The parsimony trees were built with random seeds -p 1237 and -x 1234.

The online tool iTOL (https://itol.embl.de/) was used to visualize trees.

#### Data accessibility

All raw and processed sequencing data generated in this study have been submitted to the NCBI Sequence Read Archive (SRA; https://www. ncbi. nlm. nih. gov/ sra) under accession number PRJNA1346114.

#### Cloning and Plasmid

##### pET21b-purZ-mazZ-datZ and pET21b-purZ-mazZ-datZ-dpoZ

Three genes including purZ, mazZ, and datZ from Salmonella phage PMBT28 were synthesized in three gene blocks (gblocks purchased from IDT) and were assembled (inserted) in pET21b (NdeI-XhoI). Salmonella phage PMBT28 or de novo identified phage dpoZ gene block was inserted into the XhoI site of the plasmid carrying purZ-mazZ-datZ genes as a single transcription unit from the T7 promoter.

##### pACYCT7-DNA helicase-SSB and pACYCT7-DNA helicase-SSB-dpoZ

The construct and cloning of pACYCT7-DNA helicase-SSB and pACYCT7-DNA helicase-SSB-dpoZ was described previously (Bioarchive).

##### pET28b-ldNDT

The encoding ndt gene from Lactobacillus delbrueckii subsp. Lactis DSM20072 (NCBI Reference Sequence: WP_002877839.1) were synthesized as gene blocks from IDT and were assembled into pET28b+ (NdeI-EcorI) (Javier 2018 catalysts). The resulting construct contains a N-term His-tag for protein expression and purification.

##### pET28a-NDT

The ndt gene identified from this metaGPA study were assembled into pET28a+ and the plasmids were ordered from Twist. The resulting constructs contain N-terminal His-tags for protein expression and purification.

#### NDT protein purification

ldNDT protein was expressed and purified following the protocol of a previous study (Javier 2018 catalysts). Novel identified NDT proteins were expressed in C2566 T7 express cells (NEB, catalog: C2566) and protein overexpression was induced with 0.4 mM IPTG. Cells were further grown overnight in 20 °C with constant shaking at 260 rpm. Cells were harvested via centrifugation at 5000 x g at 4 °C for 30 min. The resulting cell pellet was resuspended in a lysis buffer containing 20 mM Tris pH 7.5, 250 mM NaCl and 20 mM imidazole. Cell lysis was prepared by ultrasound sonication using a Misonix S-4000 sonicator with 20 s on and 20 s off cycles until an OD260 plateau was reached. Cell lysates were spun down at 13,000 rpm for 30 min in a pre-chilled centrifuge at 4 °C. We then combined 0.5 mL Ni-NTA agarose (Qiagen, catalog #:30210) with every 1 mL cell lysis and conducted gravity-flow chromatography. The protein bound agarose were washed twice with 5 x bed volume with cell lysis buffer. Proteins were eluted with 4 x 0.5 mL fractions in a buffer containing 750 mM imidazole, 500 mM NaCl and 20 mM Tris pH 7.5. Fractions with the protein of interest identified by SDS-PAGE were pooled and loaded onto the Amicon ultra centrifugal filter (3 kDa MWCO, Millipore, catalog #: UFC5003) to concentrate. Concentrated protein was finally recovered with a 200 ul storage buffer (20 mM Tris-HCl, pH 7.5, 100 mM NaCl, 1 mM DTT, 50% glycerol) and stored at -20 °C until its use.

#### dZ *in vivo* incorporation

Plasmids were transformed into T7 express competent cells (NEB, catalog #: C2566) according to the manufacturer’s protocol. Liquid cultures were started from single colonies and *in vivo* expression of dZ cluster genes were induced by adding 0.5 mM IPTG when OD600 reached 0.6. After overnight cell culture at 20 °C with constant shaking at 260 rpm, cell pellets were harvested by centrifugation at 13,000 x g. We then performed miniprep using Monarch plasmid miniprep kit (NEB, catalog #: T1010) to lyse cell pellets and extract plasmids. The level of dZ incorporation in these plasmids was measured and quantified by HPLC.

#### NDT reaction

The standard assay was performed by incubating 0.6 µg of purified NDT enzymes with 10 mM 2’-deoxynucleoside donor and 10 mM nucleobase acceptor in 50 mM MES pH 6.5 buffer. The reaction was incubated at 37 °C for 20 min with continuous shaking at 300 rpm. For ldNDT reactions, incubation temperature was raised to 50 °C for 5 min with continuous shaking at 300 rpm. Equal volume (40 µl) of ice-cold methanol was added into the reaction mix to stop the reaction and the enzyme was inactivated followed by heating to 100 °C for 5min. The reaction mix was diluted 1:1 with water and filtered using 0.22 µM centrifugal filters (Millipore, REF: UFC30GV00) for processing with HPLC.

#### UHPLC-MS analysis

UHPLC-MS analysis was performed using an Agilent 1290 Infinity II UHPLC equipped with G7117A Diode Array Detector and 6135 XT MS Detector, on a Waters XSelect HSS T3 XP column (2.1 × 100 mm, 2.5 µm) with a gradient mobile phase consisting of methanol and 10 mM ammonium acetate buffer (pH 4.5). The identity of each peak was confirmed by MS. The relative abundance of each nucleoside was determined by the integration of each peak at 260 nm or their respective UV absorption maxima.

## Acknowledgement

We thank Michael Kuska for providing the dZTP substrates. We thank Larry McReynolds for critical comments and discussion.

## Author Contributions Statement

WY, SX, LE designed the overall study; WY, ND, SX performed the experiments; WY and LE conducted the data analysis; WY and LE wrote the initial manuscript; SX, IRCJ and LE supervised separate part of the project; WY, SX, ND, IRCJ and LE provided critical review and feedback to revise the manuscript.

